# Inertial sensing of water content in tumor spheroids

**DOI:** 10.1101/2025.07.24.665638

**Authors:** Georgios Katsikis, Jennifer C. Yoon, Thomas R. Usherwood, Seth Malinowski, Jiaquan Yu, Chuyi Chen, Sukbom Son, Julie L. Sutton, Keith L. Ligon, Jungchul Lee, Teemu P. Miettinen, Scott R. Manalis

**Author notes:** Equal contribution (Georgios Katsikis, Jennifer C. Yoon). Drosera Biotechnologies, Cambridge, MA 02139, USA. Texas A&M College of Medicine, Dallas, TX 75246, USA.

## Abstract

Cellular water content governs the concentration of all biomolecules inside a cell, thereby influencing the physical and functional properties of the cell. However, measurements of water content in physiologically relevant cell culture models remain largely unavailable, particularly in 3D models such as tumor spheroids and organoids. Here, we achieve such measurements using a commercially available, industrial-grade, steel tube. The steel tube functions as a mechanical resonator that inertially senses the buoyant mass of particles. For microgram-scale particles ≥400 µm in diameter, we achieve <1% precision error in buoyant mass with a 5-minute acquisition interval. By sequentially measuring the buoyant mass of individual, glioblastoma patient-derived tumor spheroids in media of different densities and cell permeabilities, we determine the absolute and fractional (v/v) water content of the spheroids, along with their dry mass, volume, and density properties. We achieve ~0.4% precision error in fractional water content with a throughput of 3 spheroids per hour. This enables us to detect both inter-spheroid variability in fractional water content and acute responses to kinase inhibition. Overall, we establish a simple and accessible technique for quantifying water content in living 3D cell culture models, opening new avenues for studying biophysical regulation in multicellular systems.

## INTRODUCTION

Almost all known organisms are composed primarily of water. Within individual cells, water content defines the concentration of biomolecules, determines the extent of molecular crowding, and influences physical properties such as mechanical stiffness ^1,2^. Indeed, processes such as intracellular transport, protein synthesis, cell cycle progression, and cell divisions have all been shown to respond to, or be coupled with, changes in intracellular water content and crowding ^3–17^. These effects are relevant in all model systems, including multicellular structures. For example, stem cell self-renewal within organoids is sensitive to changes in cellular water content ^1,6^. Because water content is partially regulated by cell–matrix adhesion ^1,11,18^, mechanisms of water homeostasis may differ between isolated cells and cells embedded within multicellular environments. Yet, measurement of water content in multicellular structures remain elusive, and current research relies on osmotic perturbations of water content without direct water content quantifications. Consequently, the extent to which multicellular models display water content homeostasis and how significantly such models change their water content in response to cell state changes remains largely unexplored.

Tumor organoids and spheroids, such as neurospheres derived from brain tumors, are 3D, multicellular structures composed of cancer cells. These next generation 3D cell culture models have emerged as powerful and tractable alternatives to classical *in vitro* 2D cell lines reducing dependence on animal models. Yet, tools to measure properties and drug responses of tumor spheroids remain limited. Measuring their biophysical properties can provide valuable insights, as tumor spheroid size has been shown to influence both drug responsiveness and susceptibility to immune-mediated antitumor effects ^19–21^. More broadly, biophysical heterogeneity among tumor spheroid and organoid poses a major challenge to reproducibility in preclinical research ^22,23^. Currently, biophysical characterization of these systems relies primarily on imaging-based approaches ^22^. For example, measurements of spheroid diameter and sinking velocity have been used to estimate spheroid mass and density ^24^, and stimulated Raman scattering microscopy has been used to assess spheroid dry mass composition ^25–27^. However, the water content of tumor spheroids and other multicellular model systems has remained inaccessible to existing measurement techniques.

Here, we developed an approach for directly quantifying the water content of large (~500 μm diameter) tumor spheroids. Our approach uses an inertial resonator sensor that determines sample mass by detecting shifts in its resonant frequency induced by the presence of the sample. Such sensors have been previously applied across a wide range of biological scales, including viruses ^28^ bacteria ^29,30^, yeast ^31,32^, phytoplankton ^33^, protozoa ^34^, and mammalian cells ranging from 4-50 μm in diameter ^9,35,36^, as well as non-biological particles smaller than 50 μm in diameter ^36–38^. However, conventional resonator sensors, typically based on microelectromechanical systems (MEMS) fabricated from silicon in a cleanroom ^39^, are constrained by their small dimensions and cannot accommodate larger biological models. Conversely, larger resonator sensors constructed from glass can accommodate greater sample sizes ^40,41^ but they generally lack the sensitivity required to resolve the mass of samples in the tumor spheroid size range (300-600 μm). Our sensor bridges this gap by combining a suitable form factor for sub-millimeter biological samples with the high sensitivity necessary for accurate analysis of mass, density, and water content.

Our sensor measures the buoyant mass (*m*_*b*_) of a sample as it flows through a resonating steel tube. Buoyant mass is defined as *m*_*b*_ = *V*_*total*_(*ρ*_*total*_ – *ρ*_*fluid*_), where *V*_*total*_ is the volume of the sample, and *ρ*_*total*_ and *ρ*_*fluid*_ are the densities of the sample and the measurement fluid, respectively. By sequentially measuring the buoyant mass of the same sample in multiple fluids with distinct densities and cell permeabilities, we can determine the total volume and density ^9,32,41,42^ as well as the dry volume and density of the sample ^14,31^. This enables the quantification of the absolute and fractional water content of individual samples, such as tumor spheroids. Although similar principles have been applied to infer the average water content in populations of single cells ^17,33^, this has not previously been achieved at the level of individual 3D cell culture models. Using our method, we reveal previously inaccessible heterogeneity in fractional water content across patient-derived glioblastoma tumor spheroids and demonstrate that water content can be modulated by pharmacological perturbation. These findings illustrate the power of our approach to resolve biologically meaningful variations in water content at single-spheroid resolution.

## RESULTS

### Steel tube vibrates at higher order resonant modes

We employed an industrial-grade steel tube with a submillimeter inner diameter (*d*_*i*_ = 600 μm) as the core sensing element, serving as a mechanical resonator to measure the buoyant mass of microgram-scale particles. We selected steel for its mechanical robustness and commercial availability. Given the tube’s straight geometry and open ends, we implemented a double-clamped configuration to support vibration (Fig. 1a, S1a). Each end of the tube was screw-mounted onto a piezoelectric element with one designated for actuation and the other for sensing. The tube’s ends were connected via flexible tubing to temperature-controlled, pressurized medium vials, allowing particles to be flown back and forth by modulating the pressure between vials. A T-junction in the flow path was connected to a syringe, enabling isolation of individual spheroids within a small volume and complete exchange of the fluid inside the resonator (Fig. S1a).

**Figure 1.**
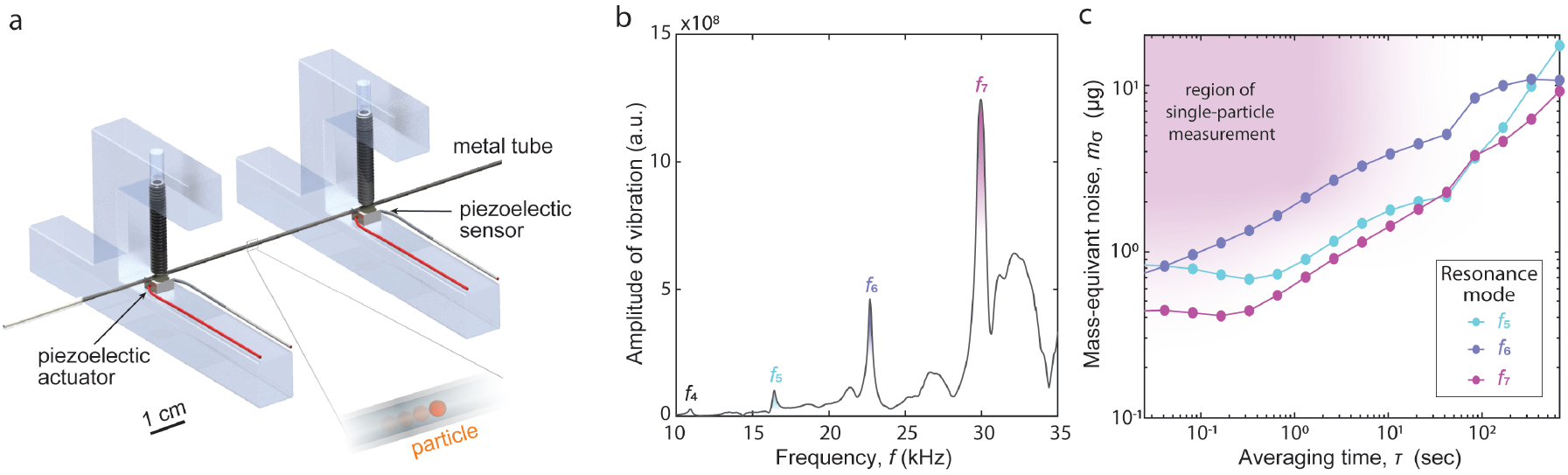
A steel tube resonator. **(a)** Schematic of a steel tube actuated to vibrate at its resonant frequencies. **(b)** Experimental amplitude of vibration vs. frequency, displaying four local maxima which correspond to the 4^th^-7^th^ modes of resonant frequencies. **(c)** Allan deviation, expressed as mass-equivalent noise, of time-series data of baseline resonant frequencies. The purple area indicates the region of measuring single particles with transit times on the order of seconds through the tube and well above the optimal noise for the 7^th^ mode.

To confirm the tube’s behavior as a resonator, we vibrated it across a range of frequencies and observed distinct amplitude peaks corresponding to its 4^th^ through 7^th^ bending resonant modes (Fig. 1b). This agreed with theoretical predictions in the kHz range (Fig. S1b). We then employed a phase-locked loop (PLL) control to selectively excite each one of these resonant modes ^43^. The 7^th^ mode exhibited the lowest mass-equivalent noise (*m*_*σ*_ ~0.5 μg), as determined via Allan deviation analysis for single-particle transits lasting several seconds (Fig. 1c). We used the 7^th^ resonant mode for all experiments from here on.

### Steel tube measures microgram-scale particles

We then assessed the performance of our system for measuring particles. Flowing 400 and 500 μm diameter polystyrene calibration beads through the vibrating steel tube yielded the expected frequency response at 7^th^ mode with phase lock loop (PLL) control (Fig. 2a) ^44^. We analyzed the frequency shifts (Δ*f*) and calibrated our measurements by calculating the effective mass, *m*_*eff*_, of the tube during the measurements of bead populations (Methods). When we separately calculated *m*_*eff*_ for the 400 μm and 500 μm beads, we observed a relative difference of 3.8%, suggesting that the calibration accuracy is within this range. Consistently, for these bead populations, our measurement-based estimates of diameter distributions matched manufacturer specifications (Fig. 2b). We then examined the precision of our system using repeat measurements of a single bead flowing back and forth through the steel tube. This revealed a coefficient of variation (CV) of 1-3% for the buoyant mass measurements, depending on bead size (Fig. 2c). Using the same data, we quantified our measurement precision as a function of acquisition time (Fig. 2d). Opting for 5 minutes of measurement (~25 repeats) yielded a precision error <1% for both 400 μm and 500 μm beads. We use a comparable acquisition time in all experiments with tumor spheroids. Finally, we examined the stability of our system during the repeated measurements of a single bead. We did not observe any correlation between the measured bead size and time (p=0.48 and p=0.49 for 400 and 500 μm beads, respectively, ANOVA). The measurement precision remained similar throughout the single bead measurement (Fig. S2a), and the measurement noise did not display any repeating patterns, as indicated by the lack of autocorrelation between consecutive bead measurements (Fig. S2b).

**Figure 2.**
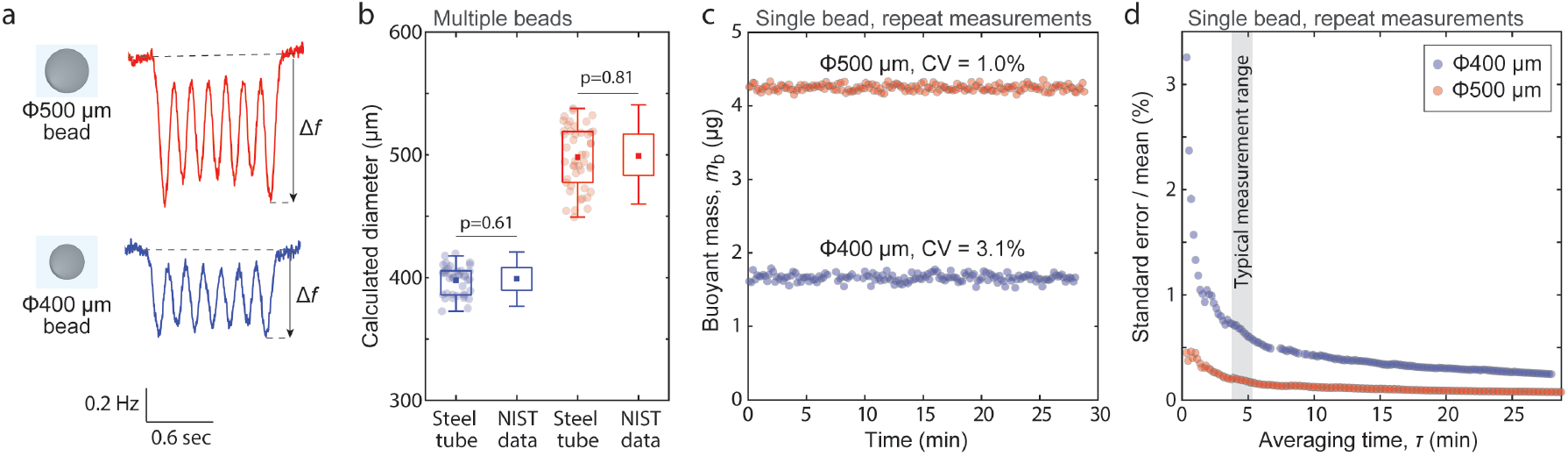
High-precision buoyant mass measurements of microgram-scale particles. **(a)** Representative signals of frequency change when flowing 500 and 400 µm polystyrene particles through the steel tube vibrating at the 7^th^ mode with PLL control (Methods). Δ*f* denotes the maximum absolute change of frequency. **(b)** Steel tube measurements of Δ*f* signals for 500 and 400 µm particles converted to diameter using the known density of particles and fluid. Particle reference size (NIST data) is shown for comparison. Individual dots depict single beads (n = 34 and 50). Reference sizes distributions are assumed normal. P-values were obtained using Student’s t-test. **(c)** Repeated Δ*f* measurements of a single particle, displayed as the buoyant mass. Data is shown for a 500 and a 400 µm particle (red and blue, respectively). n = 170 and 161 repeat measurements. CV represents the coefficient of variation. **(d)** Repeated measurement precision (standard error of mean, normalized to %) as a function of measurement acquisition time for 500 and 400 µm particles. The typical time range for repeated measurements in tumor spheroids is shown in grey.

### Quantification of absolute and fractional (v/v) water content in tumor spheroids

Having established buoyant mass measurements for microgram-scale particles, we next wanted to use our measurements to quantify the water content of individual tumor spheroids by measuring the buoyant mass of the same tumor spheroid in different fluids. First, we measured buoyant mass in normal culture medium and in high-density culture medium that contains 35% OptiPrep, which is a cell impermeable, isotonic density gradient component. With knowledge of the density of both fluids, the two buoyant mass measurements allow us to determine the total volume, mass, and density of the tumor spheroid (Fig. 3a) ^9,32,41,42^. Notably, these volume measurements are independent of the sample’s shape, thus achieving a high precision even on irregularly shaped tumor spheroids ^42^.

**Figure 3.**
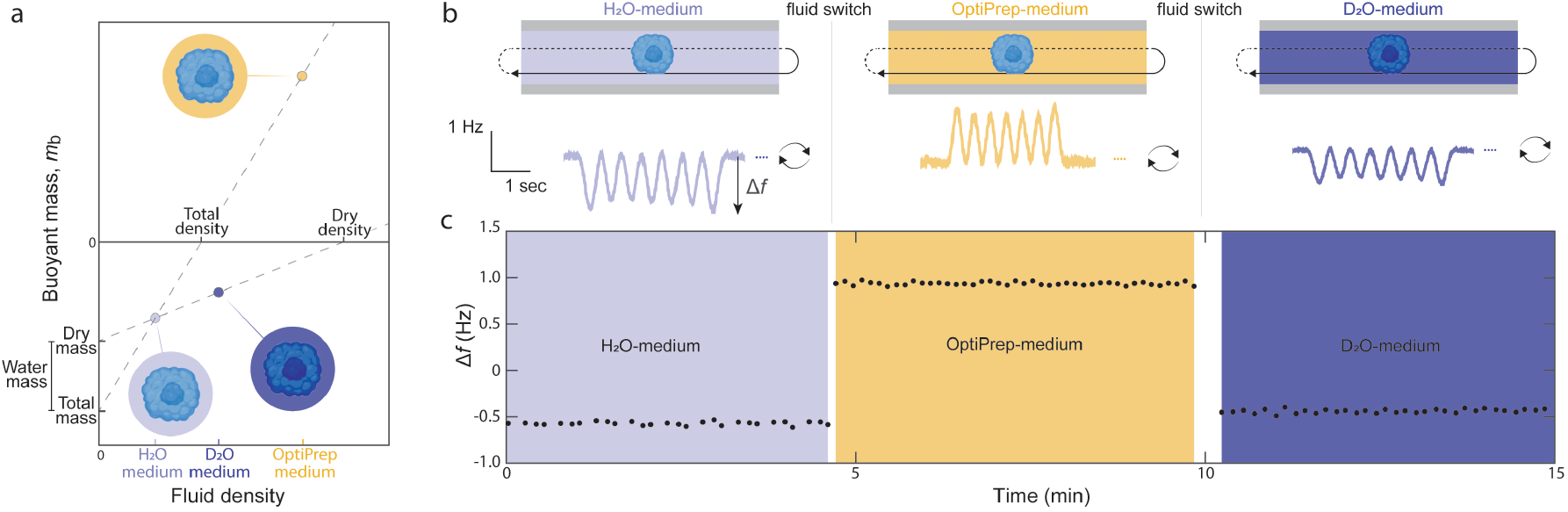
Weighing tumor spheroids in multiple fluids to determine their biophysical properties. **(a)** Schematic of our approach, which compares buoyant mass measurements in different fluids. Because water is exchanged rapidly between intracellular and extracellular environments, the H_2_O and D_2_O media (light and dark blue, respectively) have a density that corresponds to the water inside the cell. Comparing these two measurements is therefore sensitive to the dry contents of the sample. In contrast, OptiPrep-medium (yellow) is dense but cell-impermeable, and comparing measurements in H_2_O and OptiPrep-based media is sensitive to the total contents (water + dry) of the sample. **(b)** Schematic of a single tumor spheroid sequentially measured in three different fluids (top). In each fluid, the sample is flowed back and forth the resonating tube to obtain multiple buoyant mass measurements (Δ*f*) (bottom, examples of experimental signals). **(c)** Example of Δ*f* over time for a single tumor spheroid in the three measurement fluids.

Then, we measured the same tumor spheroid in medium made in heavy water, deuterium oxide (D_2_O), which has a higher density than H_2_O. However, unlike OptiPrep, D_2_O rapidly enters the cells and replaces intracellular H_2_O ^14,31,45^. As the intracellular and extracellular water now have equal densities in the H_2_O-based medium and in the D_2_O-based medium, the measurements are only sensitive to the dry components of the tumor spheroid. We can therefore consider buoyant mass as *m*_*b*_ = *V*_*dry*_ (*ρ*_*dry*_ – *ρ*_*fluid*_), where *V*_*dry*_ is the dry volume, *ρ*_*dry*_ is the density of the dry mass (i.e. dry mass / dry volume), and *ρ*_*fluid*_ is the density of the measurement fluid ^14,31^. This enables us to determine the dry volume, mass, and density of the tumor spheroid (Fig. 3a). With knowledge of the sample’s total and dry volume, we can then calculate the absolute and fractional (%, v/v) water content for individual tumor spheroids.

To implement these measurements in tumor spheroids, we utilized glioblastoma (GBM) patient-derived tumor spheroids (neurospheres) as a model system ^46^. Since OptiPrep has limited permeability to extracellular spaces within the spheroids, the measured water content reflects both intra- and extracellular water. Each spheroid was measured ~30 times over ~5 minutes in each medium (Figs. 3b&c). For medium changes, we transferred the spheroid into a syringe containing a small volume of the current medium (Fig. S1a), while the system was flushed with the new medium. Because our measurements are highly sensitive to the density of the fluid inside the steel tube, medium changes and imperfect mixing introduced fluctuations in the measurement baseline (Fig. S3a). Despite this, we obtained stable, repeatable measurements of spheroid buoyant mass (Fig. 3c), with a precision error of less than 1% in all media (Fig. S2c). Repeated measurements showed no change in spheroid mass, indicating structural integrity throughout the process. Overall, our approach enabled quantification of each spheroid’s water content within ~15 minutes. Given that brief exposure to D_2_O does not affect the viability of most cancer cells ^14^, the spheroids are expected to remain viable during the measurements.

### Detection of water content variability between individual tumor spheroids

We next sought to define water content measurement precision. This is critical for assessing the biological variability of tumor spheroids, as observed variability could arise from measurement noise alone. As there are no standard particles with fixed water content that could be used to define our measurement precision, we determined our measurement precision *in silico* by simulating buoyant mass measurements which contain similar baseline noise magnitude and slow fluctuations as the experiments (Fig. 4a, Methods). First, we determined the low frequency baseline fluctuation characteristics of experiments by fitting a 3^rd^ order polynomial to the baseline signal before and after each peak (Fig. S3b-d). We then removed the baseline fluctuations using a high-pass Butterworth filter and characterized the magnitude (expressed as standard deviation, σ) of the experimental noise (Methods). We set the water content and density properties of the simulated spheroids as equal to the experiments, with only a variation in the size to match the experiments. To ensure that our simulations were comparable to our experiments, we reanalyzed the noise magnitude in our simulated data. The noise of the simulated data corresponded to the upper (>80%) percentile of experimental noise level (Fig. 4b). Overall, these simulations allowed us to examine how noise in our measurements compound and contribute to the water content and density measurement precision.

**Figure 4.**
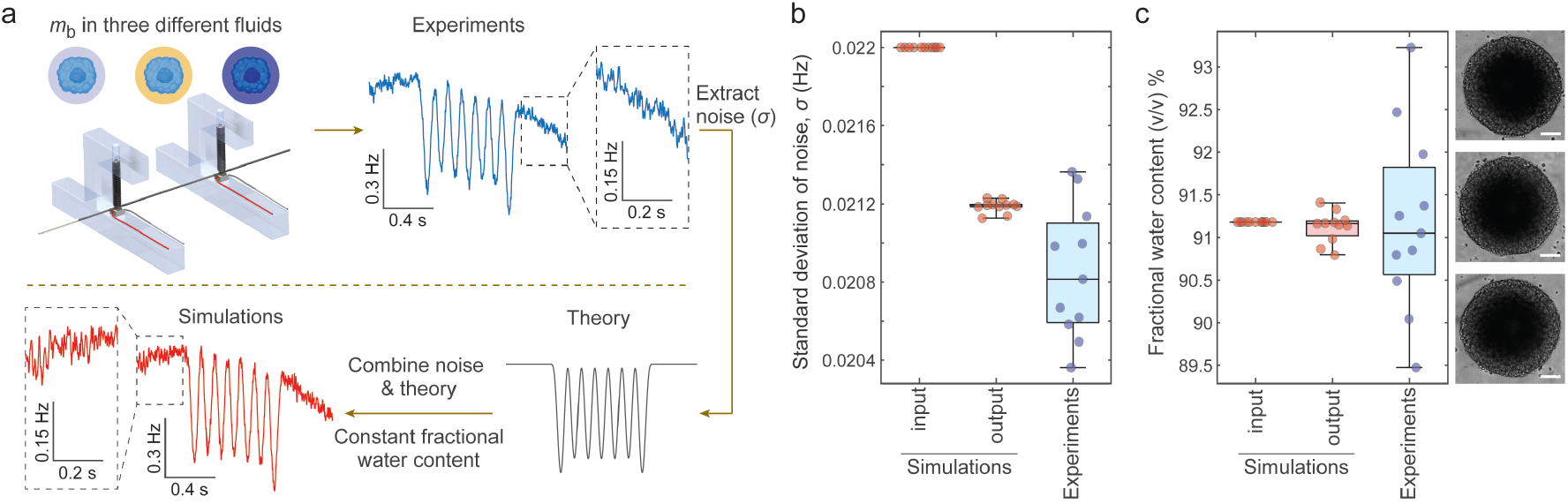
Detection of tumor spheroid water content variability. **(a)** Methodology of characterizing noise in water content measurements. The baseline noise magnitude (σ) is quantified from experiments with tumor spheroids (blue). Then, noise is simulated and added to the theoretically expected frequency shift (black) to create simulated data (red). The simulated data assumes similar variability in tumor spheroid sizes as observed in the experiments, but no variability in tumor spheroid fractional water content or density properties. **(b)** Comparison of noise magnitude between simulation input (red), output (red), and experiments (blue). Simulation output refers to the values obtained when the simulated data was analyzed identically to the experiments. **(c)** Comparison of fractional water content (v/v) between simulation input, output (red), and experiments (blue). Right displays representative brightfield images of three tumor spheroids (scalebars depict 100 µm). Each dot in the boxplots refers to a single tumor spheroid (n=11 for all conditions).

Our simulations revealed a fractional water content (v/v) measurement precision error (95% CI) of ~0.4% (Fig. 4c). The experimental tumor spheroid data showed ~6-fold more fractional water content variability than our simulations (p<0.0001, F-test of equality of variances). Thus, our measurements can resolve the water content heterogeneity between similar-sized individual tumor spheroids. Similarly, we determined that our total density measurement precision error (95% CI) is ~0.0001 g/mL and our dry density measurement precision error (95% CI) is ~0.016 g/mL (Fig. S5). Our experiments showed more variability for total and dry density, indicating that we can also resolve the heterogeneity between individual tumor spheroids.

To ensure that our fractional water content measurement precision is not biased by specific noise characteristics, we simulated tumor spheroids using different types of colored noise (slope *a* = 0 − 2 of power spectral density of noise). These simulations showed that the fractional water content precision is largely insensitive to the slope *a* (Fig. S4b). However, when the noise magnitude (σ) was increased well beyond levels observed in experiments, the noise slopes began to affect precision (Fig. S4c), highlighting the importance of maintaining high measurement quality. Additionally, we confirmed that the noise magnitude observed for each tumor spheroid did not account for the observed heterogeneity in water content (Fig. S4d).

### Staurosporine induces an acute loss of water in tumor spheroids

Having established our method for measuring water content with high precision, we next sought to examine perturbations of cellular water content. Staurosporine is a broad-spectrum protein kinase inhibitor that causes apoptosis and an acute loss of intracellular water ^42,47^. To examine the effects of staurosporine on tumor spheroids, we treated the tumor spheroids for 1 h with 2 µM staurosporine or with 0.1% (v/v) DMSO (vehicle) across two independent experiments and measured from 8-11 independent spheroids per condition. We first observed that the absolute water content of the DMSO-treated (control) spheroids varied between experiments, reflecting overall size differences of the spheroids (Fig. 5a). However, DMSO-treated spheroids in both experiments exhibited similar fractional water contents (Fig. 5b), suggesting that tumor spheroids’ fractional water content is independent of their size. In contrast, when we examined staurosporine-treated tumor spheroids, we observed significantly lower fractional water content than in the DMSO-treated spheroids (Fig. 5b). Thus, our method can detect acute changes in water content.

**Figure 5.**
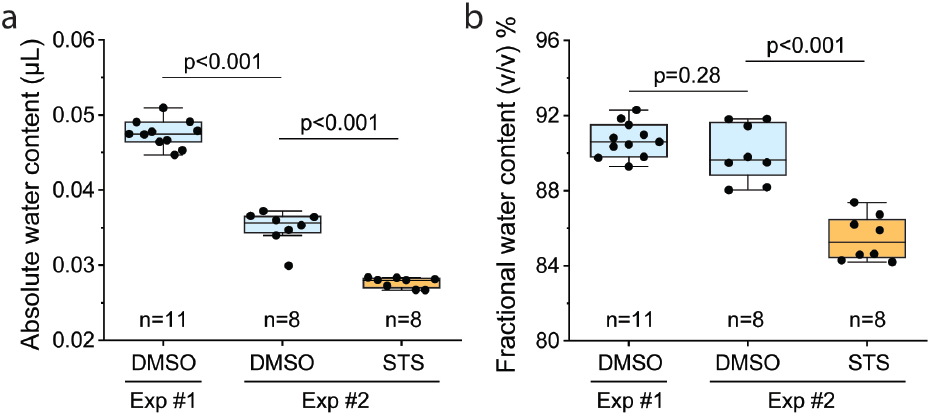
Staurosporine induces an acute loss of water in tumor spheroids. **(a)** Absolute water content (volume) and **(b)** fractional water content (%, v/v) in individual GBM tumor spheroids in two experiments. Tumor spheroids were treated 1 hour with DMSO (blue) or 2 µM staurosporine (STS, orange) prior to measurements.

## DISCUSSION

Here, we have developed an inertial sensing-based approach to quantify the water content of single tumor spheroids. This method achieves high precision (95% CI of ~0.4%), enabling us to resolve heterogeneity in water content among patient-derived glioblastoma spheroids and to detect acute changes in response to pharmacological perturbation. In addition to quantifying water content, our approach simultaneously measures the mass and density of each sample. Unlike imaging-based approaches, which often rely on assumptions about particle geometry, our approach is shape-independent and can be applied to irregularly structured biological specimens. Although demonstrated here with glioblastoma spheroids, our approach is compatible with a wide range of suspended biological particles between 300-600 µm in diameter. This range can be further extended by modifying the resonator tube dimensions. Accordingly, the method is, in principle, applicable to other tumor spheroids, organoids, very large cells (*e*.*g*., oocytes, large plankton), and even small multicellular organisms (*e*.*g*., tardigrades, *Caenorhabditis elegans*). This ability to study various 3D model systems has clinical potential given the recent emphasis of NIH and other funding bodies on the development of non-human models, especially in the context of cancer functional precision medicine ^48^.

Several limitations should be noted. First, our approach requires samples to be in suspension, which excludes adherent organoid models and matrix-embedded cultures. However, our approach is compatible with current methods using gel-supported cultures, such as alginate droplets, into solution. Second, the samples must be mechanically robust, as repeated flow-based measurements may induce shear stress that could compromise fragile structures. Third, our method relies on the use of OptiPrep to manipulate fluid density. While live cells are impermeable to OptiPrep, extracellular matrix components may permit partial penetration, potentially confounding the distinction between intracellular and extracellular water. As such, our measurements reflect total (bulk) water content, without spatial resolution between compartments. Finally, our method is relatively low throughput, with a current capacity of ~3 spheroids per hour, limiting its utility for high-throughput screening or large-scale studies.

Despite these limitations, our approach offers a unique capability: it provides quantitative, precise, single-sample measurements of water content in complex 3D biological models—something previously inaccessible with existing techniques. This opens new opportunities to investigate water homeostasis and its role in tumor biology, drug responses, and multicellular biophysics across diverse experimental systems.

## MATERIALS AND METHODS

### Fabrication and design of devices

The stainless-steel capillary tube with an outer diameter of 800 µm and inner diameter of 600 µm (#SUS304, Geumyang Materials) is placed on top of two piezo actuators (#TA0505D024W, Thorlabs) that are 5 cm apart. The capillary tubing is clamped with a setscrew that has a soft nylon tip (#AXK117, AXK). Fluorinated Ethylene Propylene tubing (#PB-0201473, IDEX Corporation) is connected at the ends of the capillary tube. The other ends of the tubing are placed inside the glass Wheaton vials and the temperature of the fluids inside the vials is maintained at approximately 37 °C with the stirring hot plate (#BZA651575, IKA).

### Operation and validation of the system

The two piezo actuators were employed where one was used for actuation and the other was used for detection. The piezoelectrically actuated signal was amplified with the custom fabricated board amplifier, while the piezoelectrically vibrated signal by the detection actuator was sent to a voltage preamplifier (#SR560, Stanford Research Systems) that uses the gain of 5,000 and the low and high-pass filter with 3 kHz and 100 kHz cut-off frequencies. The system was run in the phase-locked loop (PLL) of second order with a closed-loop mode where the resonant frequency of the 7th vibration mode was maintained with a bandwidth of 80Hz. A sampling rate of 10^8^ Hz was used with downsampling via a Cascade Integration Comb filter applying a decimation rate of 8,000. The resonant frequency signal was read with a field-programmable gate array (FPGA, #DE2-115, Terasic Inc.) connected via ethernet cable to a desktop computer. The experiments were performed using a custom code written in LabVIEW 2020 software and a computer-controlled pressure regulator (#QPV1, Proportion Air) was connected to the data acquisition (DAQ) card (#USB-6002, National Instruments) to flow fluid and particles. The frequency shift was measured when the particle was moving back and forth through the vibrating stainless-steel capillary sensor and the custom LabVIEW program reversed the direction of fluid flow when the particle was detected.

For system validation and performance analysis, National Institute of Standards and Technology (NIST) polystyrene beads with two different diameters (400 µm and 500 µm, #4340A and #4350A, Thermo Fisher Scientific) were used. For bead population studies, the magnetic stir bar was placed inside the vial containing the fluid and the beads. All beads were measured in DI water. For system cleaning following each experiment, the system was immersed in 10% v/v bleach for 5 minutes to completely wash out biological residues, after which the system was rinsed with 7% Pluronic F-127 (#P2443, Sigma-Aldrich) in DI water.

### GBM patient-derived models

Glioblastoma (GBM) patient-derived tumor spheroids have been described previously ^46^. After enzymatic and mechanical dissociation, GBM tumor resections were seeded in NeuroCult™ NS-A Basal Medium - Human (# 05750, STEMCELL Technologies) supplemented with additional growth factors, including 20 ng/mL epidermal growth factor (EGF) and 10 ng/mL basic fibroblast growth factor (bFGF) and 2 mg/mL heparin (#78006, #78003, #07980, STEMCELL Technologies). GBM tumor spheroids were passaged once every week. 1X Accutase (#AT104, Innovative Cell Technologies) was used and the tumor spheroids were incubated at 37 °C to be dissociated and single cells were resuspended in T-75 ultra-low attachment flasks (#3814, Corning) at a concentration of 2,000 cells/mL in 10 mL. The tumor spheroids would normally reach a suitable size for experiments in approximately 3 days after passage, and the spheroids were imaged using the Incucyte S3 System (Sartorius AG) to ensure appropriate spheroid diameter. Tumor spheroids were derived from the GBM model BT145. The models and more detailed information are available from the DFCI Center for Patient Derived Models (CPDM) (https://www.dana-farber.org/research/integrative-research/patient-derived-models).

### Preparation of media

35% v/v OptiPrep medium was made by mixing OptiPrep density gradient medium (#92339-11-2, Sigma-Aldrich) with Neurocult medium. 1x PBS with heavy water was made by adding 10% v/v 10x PBS (#AM9624, Invitrogen) with 90% v/v deuterium oxide (#151882, Sigma-Aldrich). The media were vacuum filtered using the filter unit (#73520-990, VWR International) and their densities were measured using the Mettler Toledo Precision Balance (#ME103TE, Hogentogler & Co.) as *ρ*_*H*2*0*−*medium*_= 1037.8. *ρ*_*optiperp*−*medium*_= 1141.8. *ρ*_*D*2*0*−*medium*_= 1131.3 kg/m3, respectively for water-based neurocult medium, neurocult medium with 35% OptiPrep, and D_2_O-based PBS.

### Staurosporine treatment of GBM tumor spheroids

For staurosporine measurement, GBM BT145 cells were seeded in 96-well round-bottom ultra-low attachment plates (7007, Corning Inc., Corning, NY, USA) at a concentration of 6,000 cells in a volume of 90 µL. After sufficient tumor spheroid formation, the tumor spheroids were exposed to 2 µM staurosporine (Abcam) or a control (0.1% v/v DMSO) for 1 hour. The tumor spheroids were measured immediately after drug exposure.

### Analysis of frequency responses

The raw data were processed using MATLAB 2019b and the Signal Processing Toolbox was utilized. Savitzky-Golay filter was applied to the resonant frequency measurements from the sensor. The resulting shift Δ*f* in resonant frequency due to a particle passing through the steel tube vibrating at the 7^th^ resonant mode consists of seven local minima (for H_2_O and D_2_O) or maxima (for OptiPrep) altogether called antinodes, which we collectively refer to as a peak. In our signals, both from experiments and simulations, the peaks were detected by finding antinodes with prominence around 5-10 times higher than the standard deviation of noise. The antinodes were identified and grouped together in sets of seven that constitute a single peak. For a given peak, its baseline resonant frequency *f* was defined by fitting a 3^rd^ degree polynomial before and after the peak, taking for both three times the interval between the first and second antinode. Only the extreme (first and last) antinodes were used for buoyant mass determination because they have the maximum absolute shift |Δ*f*| in resonant frequency. To smooth out the distortion of the antinodes due to noise, we fitted local 2^nd^ degree polynomials to the antinodes and took the local maximum or minimum. A given peak was accepted only if the two antinode signals have a maximum of 20% relative difference. Then, the height of each peak Δ*f* was converted to buoyant mass by

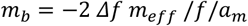

where *a*_*m*_ is the mass discrepancy ^49^. The calibration of the steel tube was done by calculating *m*_*eff*_ via known buoyant masses of experimental measurements of NIST polystyrene beads, assuming mean diameters from vendor. For the spheroid measurements, the effective mass *m*_*eff*_ was adjusted accounting for the densities of the different fluids used.

### Determining water content

We determined cellular water content from buoyant mass measurements according to a previously established approach ^14,17,31^. In short, buoyant mass is defined as

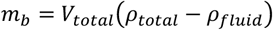

where *m*_*b*_ is buoyant mass, *V*_*total*_ is total volume, *ρ*_*total*_ is the buoyant density of the tumor spheroid, and *ρ*_*fluid*_is the density of the measurement fluid. We determined the total volume of the tumor spheroids by comparing buoyant mass measurements in normal medium and OptiPrep-based medium using to the equation above.

We determined the dry volume of the tumor spheroid by separating the buoyant mass contribution of tumor spheroid’s dry content and water content.

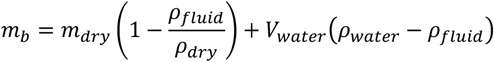

where *m*_*dry*_ is the dry mass, *ρ*_*dry*_ is the density of dry mass, *V*_*water*_ is the volume of water in the sample, and *ρ*_*water*_ is the density of water. The first term of the equation depicts the dry content contribution to *m*_*b*_, and the second term depicts the water content contribution to *m*_*b*_. When comparing tumor spheroid buoyant mass measurements in normal medium and D_2_O-based medium, the latter term of the equation approximates to zero, allowing us to solve for the first term. The absolute and fractional water content (v/v) were calculated based on the dry and total volumes.

### Determination of noise in experimental signals

The segments of the data between peaks were isolated. To characterize noise, first the low frequency (*f* < 1) drift was removed by applying a Butterworth high-pass signal *f*_*high*−*pass*_ = 1 Hz and order *n* = 2. The mean of the segment was subtracted from the segment and the standard deviation σ of the noise was calculated in the time domain. In addition, the slow drift of the frequency baseline was characterized via the parameters of the 3^rd^ degree polynomial fits applied to immediate baselines before and after each peak, taking for both three times the interval between the first and second antinode. For the characterization of the slow baseline drift, extreme outliers were excluded.

### Simulations of noise and frequency responses

The simulations were performed by taking the theoretical formula Δ*f* for frequency change at 7^th^ resonant mode and adding color noise with standard deviation σ and slope *a* (*a* = 0 − 2) of the form *S* ∝ 1/*f*^*a*^, where *SS* is the power spectral density of the noise. For the simulations, we used σ values in the upper (>80%) percentile of the noise level observed in the experiments with tumor spheroids, and we assumed white noise (*a* = 0). We selected these values to confidently determine an upper limit to the precision of our water content measurement. In addition, a slow baseline drift was applied to simulated signals. This drift was determined from experiments by taking normal distributions for the mean and standard deviation of the parameters of the 3^rd^ degree polynomial fits applied to the baselines before and after each peak in experiments with tumor spheroids. Since the experiments are produced by applying a PLL of 2^nd^ order to excite the tube at the 7^th^ resonant mode, the experimental signal can be simplified to a first-order Butterworth low-pass filters applied to the peak and the noise ^43^. Therefore, an equivalent low-pass filter with cut-off frequency equal to the bandwidth of the PPL loop (80 Hz) was applied to the simulations. Data (frequency responses) were simulated for a similar number of repeat buoyant mass measurements as in the experiments, assuming the same fluid densities and the same number of tumor spheroids as in the experiments. The data from the simulations were analyzed identically to the data from the experiments.

### Statistics and data presentation

Statistical comparisons were carried out using Welch’s t-test, unless otherwise stated. All p-values were calculated using OriginPro 2025 software. In all box plots, the central line and square depict the median and mean, respectively, the bottom and top edges of the boxes respectively indicate the 25th and 75th percentiles, and the whiskers indicate the 5^th^ and 95^th^ percentiles. When full data was not available (manufacturer provided bead reference data), percentiles are defined by assuming data follows a normal distribution.

## ACKNOWLEDGEMENTS

This work was supported by the D.K. Ludwig Fund for Cancer Research (S.R.M.), the MIT Center for Precision Cancer Medicine (S.R.M), the Koch Institute Support (core) Grant P30-CA014051 from the National Cancer Institute, the NIH’s National Institute of General Medical Sciences (Award R01GM150901) to T.P.M. and S.R.M., and the National Research Foundation of Korea (NRF) grant funded by the Korea government (MSIT) (RS-2025-00560250) to J.L.

## Author contributions

G.K., J.L., T.P.M., and S.R.M. designed the research; S.S. prepared and provided the steel tube resonator; J.C.Y. built the measurement setup with guidance from J.L.S. and G.K.; J.C.Y and C.C. carried out the experiments with assistance from G.K.; S.M., J.Y., and C.C. carried out tumor spheroid culturing; G.K. and T.R.U. analyzed the data and carried out the simulations; K.L.L., and S.R.M. contributed resources; G.K., T.P.M., and S.R.M. supervised the work and wrote the paper with input from K.L.L. All authors approved the manuscript.

