## Supplemental Figures for "Inertial sensing of water content in tumor spheroids"

**Affiliations:**
[1] Koch Institute for Integrative Cancer Research, Massachusetts Institute of Technology, Cambridge, MA 02139, USA.
[2] Harvard-MIT Department of Health Sciences and Technology, Institute for Medical Engineering and Science, Massachusetts Institute of Technology, Cambridge, MA 02139, USA.
[3] Department of Pathology, Dana-Farber Cancer Institute, Harvard Medical School, Boston, MA 02115, USA
[4] Department of Mechanical Engineering, Korea Advanced Institute of Science and Technology, 291 Daehak-ro, Yuseong-gu, Daejeon 34141, South Korea
[5] Center for Patient-Derived Models, Dana-Farber Cancer Institute, Boston, MA 02115, USA
[6] Department of Pathology, Brigham and Women's Hospital, Harvard Medical School, Boston, MA 02115, USA
[7] Department of Mechanical Engineering, Massachusetts Institute of Technology, Cambridge, MA 02139, USA.
[8] Department of Biological Engineering, Massachusetts Institute of Technology, Cambridge, MA 02139, USA.

[*] Equal contribution
^ Present address: Drosera Biotechnologies, Cambridge, MA 02139, USA
+ Present address: Texas A&M College of Medicine, Dallas, TX 75246, USA

**This file includes:**
Supplementary Figures S1 – S5

### SUPPLEMENTARY FIGURES

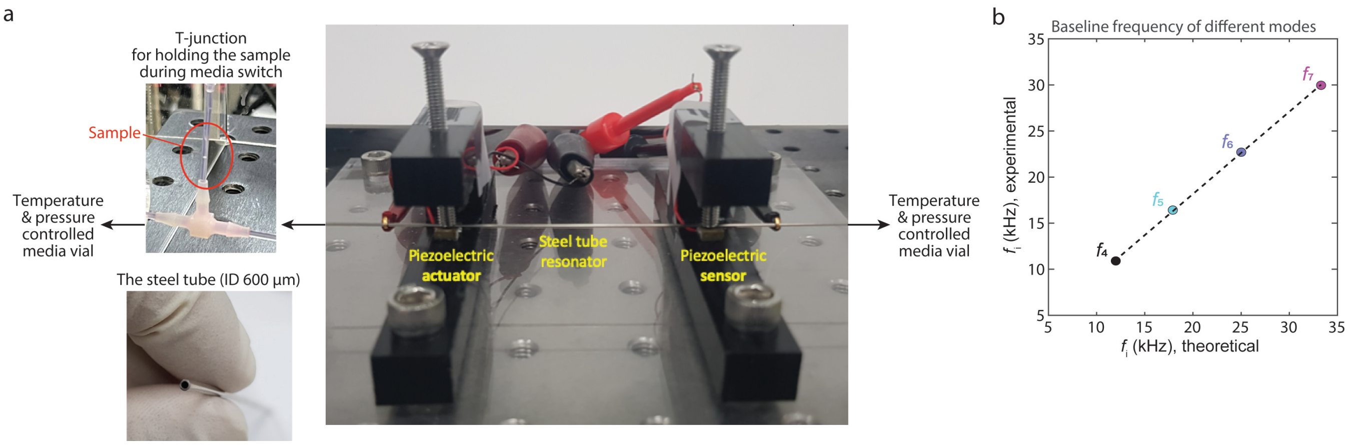


**Figure S1. Details of the steel tube-based measurement setup**

**(a)** Images of the steel tube-based measurement setup. At each end, the steel tube is connected to tubing leading to temperature and pressure-controlled medium vials. These pressure controls are used to adjust the fluid flow in the steel tube. On one side, the connecting tubing contains a T-junction with a syringe, which can be used to temporarily hold the measured sample in a small and separate volume, as the measurement fluids in the source vials and the steel tube are changed. The bottom left displays an image of the steel tube when held separately. **(b)** Correlation between the theoretical and experimental frequencies of each resonant mode.

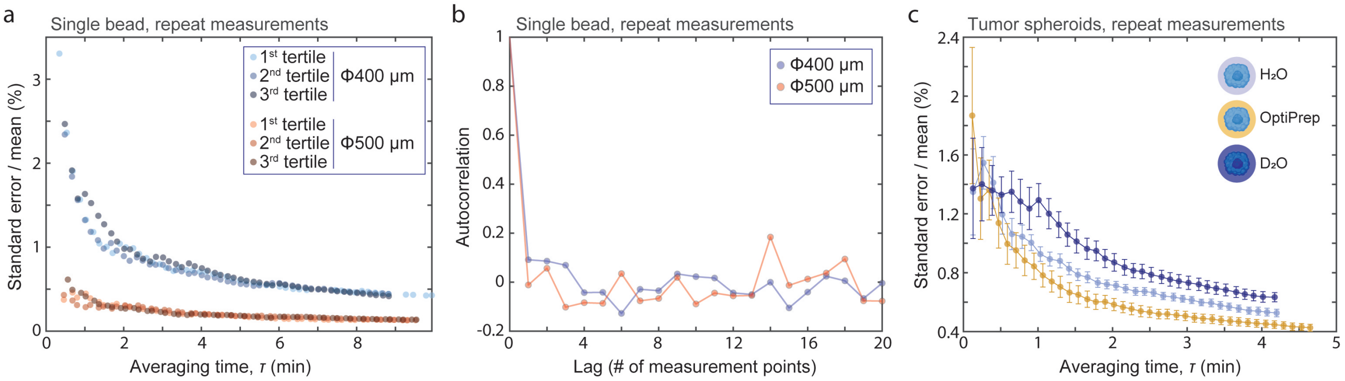


**Figure S2. Characterization of buoyant mass measurement precision**

**(a)** Repeated measurement precision (standard error of mean, normalized to %) as a function of measurement acquisition time for 500 and 400 µm polystyrene particles. Data is the same as in Fig. 2d, but each trace is separated into three sections to exemplify the stability of the measurements. **(b)** Autocorrelations of repeated measurements of 500 and 400 µm polystyrene particles. **(c)** Repeated measurement precision (standard error of mean, normalized to %) as a function of measurement acquisition time for GBM tumor spheroids in three different fluids. Data depicts mean and SEM of 11 separate tumor spheroids.

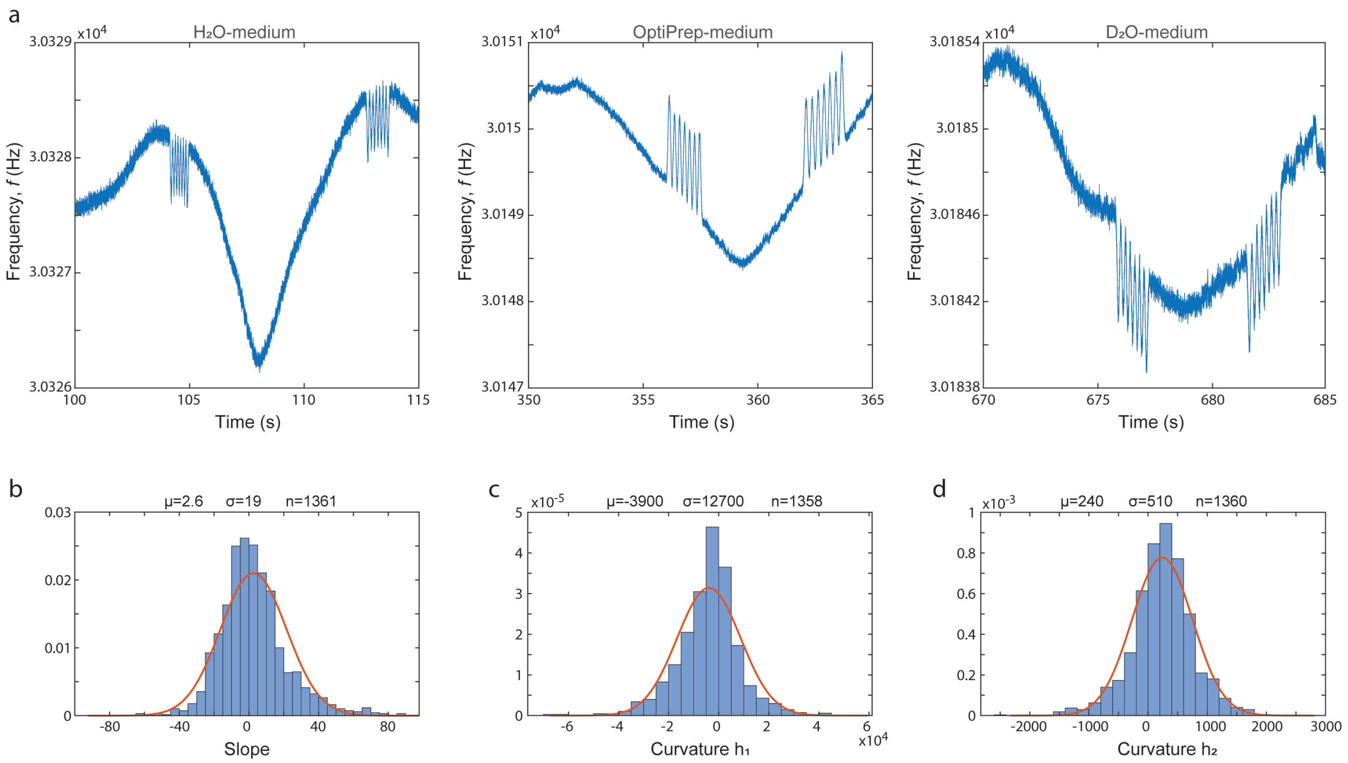


**Figure S3. Buoyant mass measurements in multiple media contain baseline fluctuations.**
**(a)** Representative raw data of GBM tumor spheroid measurements in three different media. As a small volume of the fluids is carried over when the fluids are changed, the incomplete mixing of the fluids results in baseline fluctuations. Note that the y-axis values are relative to a reference value and cannot be compared between plots. **(b-d)** Quantification of the low frequency ($f < 1\ Hz$) drift of the baseline. Each histogram depicts a separate parameter of the 3rd degree polynomial fits applied to immediate baselines before and after each peak. Curvature $h_1$ refers to the 2nd order curvature, and curvature $h_2$ refers to the 3rd order curvature. $\mu, \sigma, n$ refer to mean, standard deviation, and number of data points, respectively.

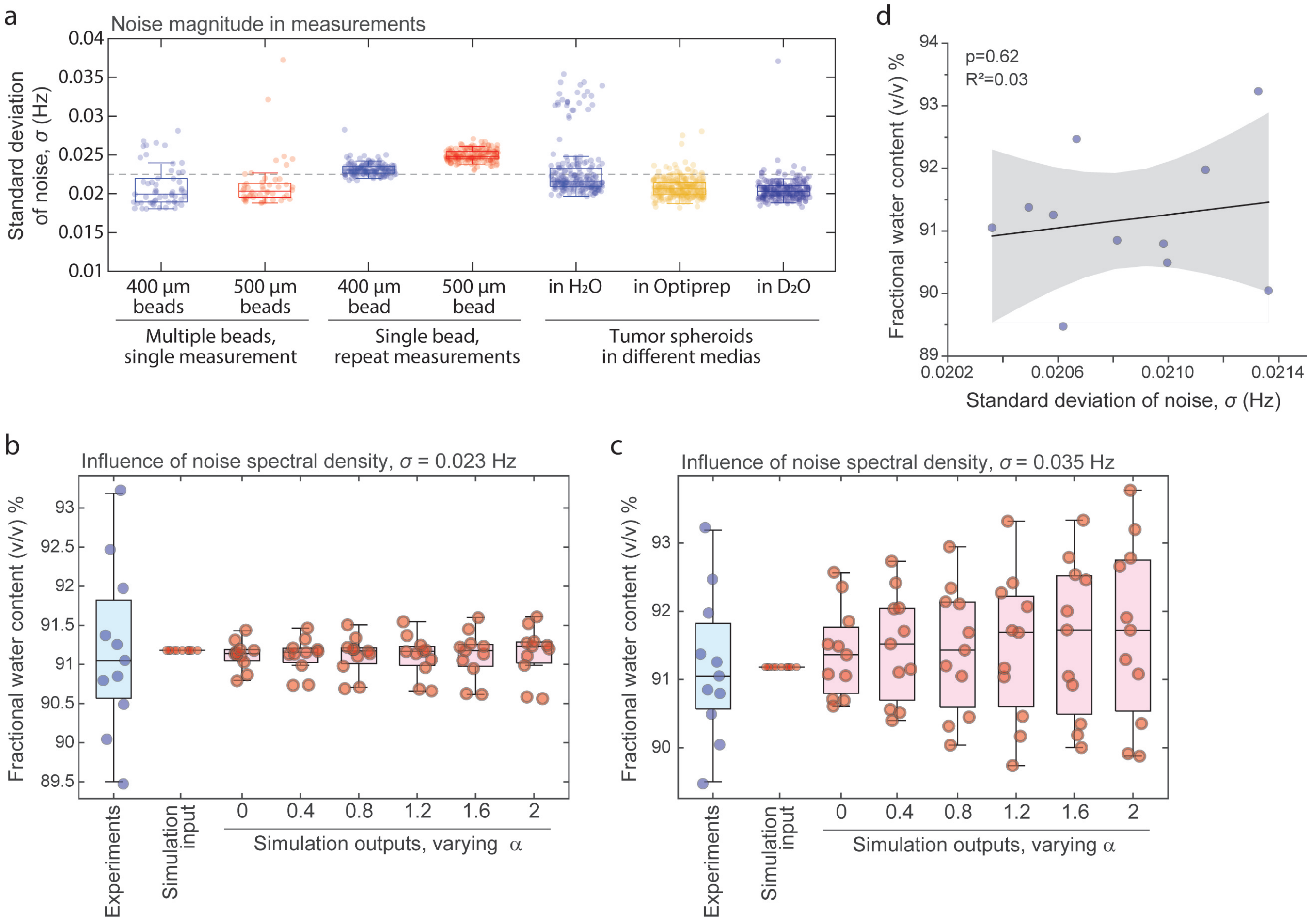


**Figure S4. Evaluation of water content measurement noise in experimental and simulated datasets.** **(a)** Baseline noise magnitude ($\sigma$) from indicated experiments/experimental conditions. On average across all data, we observed that $\sigma$~0.0225 Hz (dashed horizontal line). Each opaque dot depicts a baseline section between two buoyant mass measurements. **(b-c)** Fractional water content in experimental and simulated data. Simulations were run with indicated noise spectral density slope values ($\alpha$) using either low (b) or high (c) noise magnitude ($\sigma$). The lower noise magnitude (panel b) is similar to the noise observed in tumor spheroid experiments. Each dot depicts a separate tumor spheroid (n=11). **(d)** Correlation between baseline noise magnitude ($\sigma$) and the derived fractional water content in experiments. Each dot depicts a separate tumor spheroid (n=11). Black line and shaded area depict a linear fit and 95% confidence bands. Correlation p-value was obtained using ANOVA. The correlation was statistically insignificant (p=0.62) also when analyzed from simulated data.

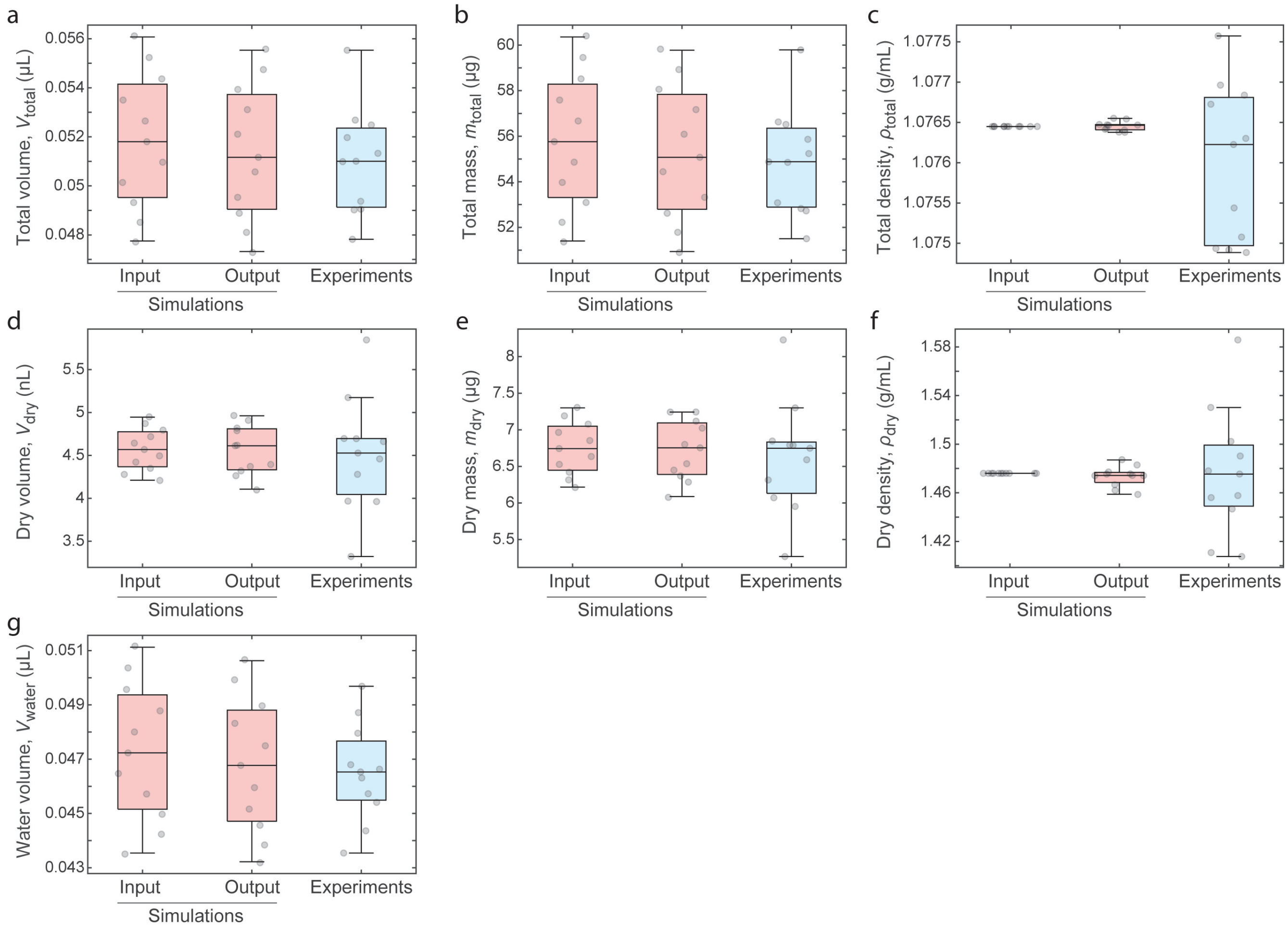


**Figure S5. Simulated and measured biophysical features of GBM tumor spheroids.**
**(a-g)** Simulated (input and output, red) and experimentally determined (blue) GBM tumor spheroid total volumes (a), total masses (b), total densities (c), dry volumes (d), dry masses (e), dry densities (f), and water volumes (g). Each opaque dot depicts a separate tumor spheroid (n=11).
